# Environmental feedback maintains cooperation in viruses

**DOI:** 10.64898/2026.04.20.719708

**Authors:** Jaye Sudweeks, Christoph Hauert

## Abstract

Cooperators that pay a cost to provide benefits to others are vulnerable to exploitation by defectors that reap the benefits while avoiding the costs. Thus selection on the individual level can lead to loss of cooperators while lowering overall population fitness. Accordingly, the maintenance of cooperation is a key problem in evolutionary biology. The puzzle of cooperation extends to viruses: cooperative viruses produce gene products that can be shared, while defector viruses produce less and instead use products made by cooperators. In coinfection, defectors are always advantaged, predicting the loss of cooperation. However, the fitness of cooperator and defector phenotypes is context dependent. Though defectors are advantaged in coinfection, they suffer reduced replication in single infections. Environmental feedback is the process by which changes in population composition alter viral densities and rates of coinfection, which in turn feed back to affect the fitness of each type. We show that environmental feedback maintains cooperation in viruses. We also find that defector emergence may interfere with phage therapy by disrupting the phage dynamics that cause host extinction, and demonstrate that the introduction of defectors lowers viral densities and drives viral extinction, suggesting that defectors that replicate alone could function as antiviral therapies.

## 1 Introduction

Cooperators that provide a benefit to others at a cost to themselves are vulnerable to exploitation. Because natural selection operates at the level of the individual and increases the abundance of the fittest type, defectors that forgo a cost but still receive a benefit should prevail, and cooperation should be rare. Using the tools of evolutionary game theory (EGT), which applies concepts from game theory to populations evolving under natural selection [1], we can represent this tension formally through the payoff matrix in Eq. (1),

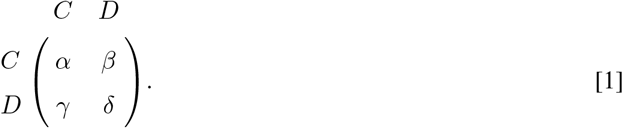

Rows correspond to the phenotype of the focal individual, columns the phenotype of interaction partner, and each entry the payoff that the focal individual receives after an interaction. Accordingly, a cooperator (C) interacting with another cooperator receives payoff *α*, a cooperator interacting with a defector (D) receives payoff *β* while the defector receives payoff *γ*, and a defector interacting with another defector receives payoff *δ*. The ordering of the payoffs determines the characteristics of the interaction. The ordering *γ* > *α* > *δ* > *β* is the prisoner’s dilemma: defectors receive a higher payoff than cooperators regardless of partner type (*γ* > *α* and *δ* > *β*) and should spread to fixation, though this lowers population fitness (*δ* < *α*).

Despite its apparent vulnerability, cooperation is abundant in the natural world [2–5]. Even viruses exhibit a wide array of cooperative traits [6] including the production of shared intracellular goods. During infection, viruses produce essential gene products in the host cell environment. Coinfection by multiple viruses is possible, so if a virus cannot guarantee exclusive access to its own gene products they may be shared by all viruses that coinfect the cell. A number of viral proteins are known to be shared, including replicase proteins, which replicate viral genetic information, and capsid proteins, which comprise the viral capsid that transports viral progeny between cells; indeed, a given viral genome may code for multiple shared goods [7]. Cooperator viruses contribute to the pool of shared goods, while defectors emerge through mutations that remove genes coding for the shared product, so that defectors replicate using products produced by the cooperator. Such mutations are typically deletions that make defector genomes shorter, enabling faster replication [8, 9]. In bacteriophage *ϕ*_6_, a virus of bacteria, the payoff ordering (in terms of replicative output after coinfection) corresponds to a prisoner’s dilemma [10]: defectors should spread to fixation, decreasing population fitness or even causing viral extinction [8, 11]. However, the cooperative *ϕ*_6_ phenotype persists.

Faced with a contradiction between expectation and observation, identifying mechanisms that maintain cooperation is a fundamental problem in evolutionary biology [12]. Previous work in EGT has suggested a number of mechanisms that maintain cooperation [13–21]. Much work assumes that all fitness differences between two phenotypes arise from interactions between the phenotypes. In the viral setting, this assumption is violated: viruses can infect and replicate in cells alone, introducing fitness effects independent of any interaction. This means the standard predictions— for example, that under a prisoner’s dilemma defection should spread to fixation— need not hold, and ecological context becomes critical [22].

We propose that environmental feedback maintains cooperation by shifting the relative fitness of cooperators and defectors. We define the environment as anything outside the individual that impacts its fitness [23, 24]. Environmental feedback is the process by which changes in population composition alter the environment, here viral densities and corresponding coinfection rates, which in turn feeds back to affect the fitness of each type. The fitness of viruses is inherently context-dependent: cooperators have high fitness when infecting a cell alone, while defectors experience reduced replication in the absence of cooperators [25–27]. The dynamics change fundamentally under coinfection, where defectors have elevated replication at the expense of cooperators. Crucially, rates of coinfection are not static but depend on viral density [28, 29]: when density is high, coinfection is common and defectors benefit from cooperator products; when density is low, single infections predominate and defector fitness declines.

To see how this feedback might unfold, consider a cooperator-only population at high density. Rare defectors have a replicative advantage because coinfection is common and defectors will coinfect with cooperators that they can exploit. Defectors increase in abundance. However, as defectors become more common, the likelihood of defector-defector coinfection increases. Coinfecting defectors have low rates of replication, so viral density falls, and single infections become more frequent. Defectors still experience reduced rates of replication, but in contrast cooperator replication is unimpeded, and cooperators rebound. Environmental feedback may lead to oscillations in cooperator and defector densities, or the effects of single infections and coinfections may balance, resulting in stable coexistence of the two types at mixed equilibrium.

We develop and analyze an ODE model, inspired by bacteriophages, which explicitly tracks the densities of viruses and hosts, allowing the rates of single versus coinfection to vary. Viral defectors range from full defectors [8, 9, 30, 31], which produce no shared good and cannot replicate without a cooperator, to partial defectors [10, 27], which produce less of the shared good than cooperators but can still replicate alone [25]; we focus on partial defectors. We find regimes where environmental feedback alleviates the prisoner’s dilemma between cooperators and defectors, leading to either coexistence or fixation of the cooperators. However, we also find regimes where environmental feedback is not sufficient, and cooperation is lost, leading to either fixation of defectors or extinction of the viral population overall.

## 2 Model derivation

Our model is inspired by the dynamics of bacteriophages and their bacterial hosts. In the absence of viruses, uninfected hosts (*H*_*U*_) undergo logistic growth with net per-capita growth rate *r* and competition rate *ξ*. Cooperator (V_*C*_) and defector (V_*D*_) viruses infect uninfected hosts. Single infections occur at per-capita rate *µ*. Empirical work finds that coinfection is often limited to few viruses per cell [10]; we set the limit at two viruses per host for mathematical convenience. We assume that multiple infections are only possible in a very small time window, so singly infected host cells cannot be infected again; it has been shown that given sufficient time many viruses can prevent multiple, sequential infections through host cell manipulations [32–34]. Simultaneous double infections occur at per-capita rate *µ*_2_. It is natural to assume that each infection in a double-infection event occurs at per-capita rate *µ*, so that *µ*_2_ = *µ*^2^. Infected hosts lyse (rupture) at per-capita rate *d*, releasing viral replicates. Because lytic infection rapidly redirects host cellular machinery toward viral replication [35, 36], we assume that infected hosts are passive and do not compete with uninfected hosts for resources for replication.

The number of replicates produced by each infecting virus depends on its own strategy and that of its coinfecting partner, if any. Let us first focus on the replicates produced by coinfecting viruses. We assume that cooperators carry multiple genes coding for shared products. Defectors emerge through mutations that delete part of one or more of those genes. We assume that defectors can still produce the shared goods, though in far smaller quantities than cooperators, but their shorter genome allows for faster replication. Cooperators pay a cost *c* in replication speed to provide benefit *b* to coinfecting viruses through shared goods production. The payoff matrix in Eq. (2) describes the relative rates of replication for each type after coinfection:

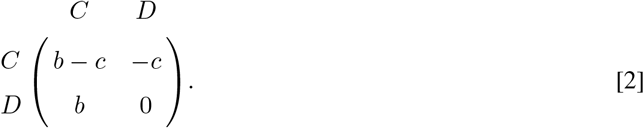

The replication advantage that defectors gain by shortening their genome is independent of the phenotype of any coinfecting virus. We can represent this advantage by adding *c* to each entry, which does not alter the evolutionary dynamics.

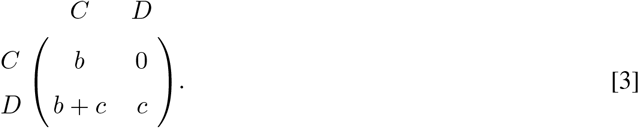

Finally, dividing by *b*, which again does not alter evolutionary dynamics, reduces the matrix to the single parameter *ρ* = *c*/*b*, setting cooperators as the reference type with relative replication one,

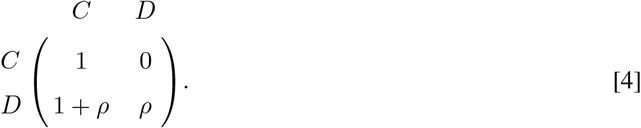

The parameter *ρ* is the ratio of the cost cooperators pay to the benefit they provide. Equivalently, *ρ* is the temptation to cheat, or the relative replication advantage defectors gain because of their shorter genomes. Because this advantage derives from genome length rather than interactions, *ρ* appears in all defector payoffs: as a boost in coinfection with cooperators (1 + *ρ*), and baseline replication when coinfecting with other defectors (*ρ*). In our analysis, we vary *ρ*: when *ρ* is small, the benefits of receiving the shared product far outweigh the advantage gained through a shorter genome, while for *ρ* → 1 the benefit gained by defecting is almost equal to the benefit of cooperation. Note that we require 0 < *ρ* < 1; for *ρ* = 0 defectors are identical to cooperators, while for *ρ* ≥ 1 two coinfecting defectors have higher payoffs than two coinfecting cooperators and viruses no longer face a prisoner’s dilemma, which requires that mutual cooperation is more productive than mutual defection. The donation game provides an intuitive, single parameter description of the prisoner’s dilemma that viruses face in coinfection but is restricted to partial defectors and cannot capture full defectors that cannot replicate without cooperators.

The relative rates of replication shown in Eq. (4) are scaled by the average burst size, λ. For example, a host cell infected by two defectors releases 2λ*ρ* defector replicates. We assume lysis occurs once host resources are exhausted and that the viral replication rate is high. Consequently, lysis time and the total count of viral particles released are independent of whether one or two viruses of the same type infected the cell. Accordingly, we can combine single and double infection events of a single type into the same infected host class (*H*_*C*_ or *H*_*D*_). Note that this means that *ρ* also determines a defector’s productivity in single infection.

We also assume that viral densities are sufficiently higher than hosts such that impacts of infection events on the viral pool can be neglected. Viruses degrade at the rate *dκ*; for *κ* > 1 viruses are shorter lived than infected hosts, while for *κ* < 1 viruses outlive infected hosts. Together, these processes yield Eq. (5).

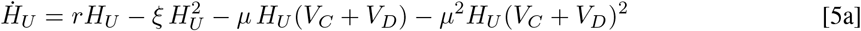

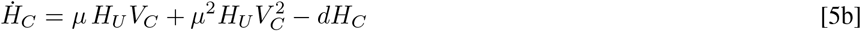

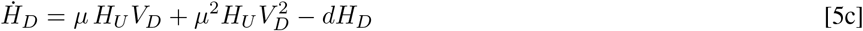

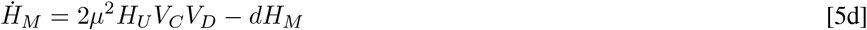

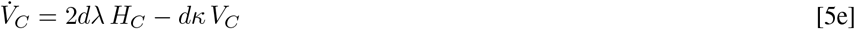

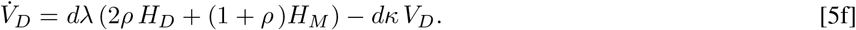

It is worth noting that selection acts on viruses in host cells, where cooperators and defectors replicate. In coinfected cells, Eq. (4) determines relative rates of replication; if defectors and cooperators coinfect, defectors exploit cooperator-produced goods and replicate at a higher rate. In singly infected cells, each type replicates independently at its intrinsic rate, with cooperators outperforming defectors. Environmental feedback determines how often viruses encounter each context. The terms *µH*_*U*_(*V*_*C*_ + *V*_*D*_) and *µ*^2^*H*_*U*_(*V*_*C*_ + *V*_*D*_)^2^ describe the rates of single and double infection, respectively: single infection depends linearly on total viral density while double infection depends on its square. When viral density is high, coinfection is common and defectors are advantaged; when viral density is low, single infections dominate and cooperators are advantaged. Note that the 2 in Eq. (5d) is a combinatorial term that results from (*V*_*C*_ + *V*_*D*_)^2^.

## 3 Results

To investigate the impact of environmental feedback on cooperation in viruses, we conduct bifurcation analysis using *ρ* as the bifurcation parameter. To build intuition, we first highlight important underlying structure from analysis of the system with a single viral type [37]. We then consider the full system with both viral types, treating our equations analytically as far as possible before turning to numerical analysis for three parameter regimes of interest. We consider the implications for the fate of cooperation as well as two viral applications: phage therapy and antiviral therapy using defectors.

### 3.1 Dynamics of a single viral type

To understand the dynamics of Eq. (5), it is useful to first consider a population of only cooperator or defector viruses. The corresponding single-type system [37] admits five equilibria: the extinction equilibrium (*E*_0_, no hosts or virus), the uninfected equilibrium (*E*_*U*_ = *r*/*ξ*, only uninfected hosts), and three infected equilibria (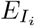, hosts and virus), of which at most two are biologically relevant. A key structural result is that the number of biologically relevant infected equilibria depends on r relative to *µ*^2^/*µ*_2_ = 1: below this threshold there is one; above it there are up to two.

### 3.2 Equilibria and stability

Equation (5) admits up to nine equilibria of the form (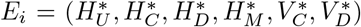), labeled by index *i*. In each, 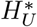 refers to the equilibrium density of uninfected hosts, 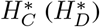 to the equilibrium density of hosts infected by only cooperator (defector) viruses, 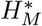 to the equilibrium density of hosts coinfected by both a cooperator and defector, and 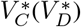 to the equilibrium density of cooperator (defector) viruses in the environment. Eight equilibria we encountered in the previous section: the extinction equilibria *E*_0_, the uninfected equilibrium *E*_*U*_, three cooperator equilibria *E*_*Ci*_, and three defector equilibria *E*_*Di*_. In addition, there may be a single mixed equilibrium *E*_*M*_ .

From Eq. (5a) we see that *E*_0_ is unstable whenever *r* > 0 and hosts are able to grow. To determine the stability of all other equilibria, we calculate the Jacobian of our system, see appendix S1. The uninfected equilibrium *E*_*U*_ is unstable when either λ > λ_*T*_ = (*dξκ*)/(2*µr*) or *ρ* > *ρT*_*DU*_ = (d*ξκ*)/(2λ*µ*r). The stability conditions for *E*_*U*_ have a natural biological interpretation in terms of the basic reproductive number, *R*_0_, of each viral type: the per capita rate of infection is *µ* and viruses have an expected lifespan of 1/(*dκ*). A cooperator introduced into an entirely uninfected host cell population at its equilibrium (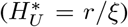) therefore infects (*rµ*)/(*dκξ*) cells on average. Note that double infections are negligible when phage densities are small. An infected host has an expected lifespan 1/*d* and produces new cooperator viruses at rate 2*d*λ. Overall, each cooperator initially produces *R*_0_ = (2λ*rµ*)/(*dκξ*) new viruses, and cooperators can invade if *R*_0_ > 1. Solving *R*_0_ = (2λ*rµ*)/(*dκξ*) > 1 for λ yields the condition above. An analogous derivation follows for defectors.

Stability conditions for 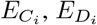, and *E*_*M*_ are analytically intractable, so we turn to numerical stability analysis for illustrative sets of parameters.

### 3.3 Bifurcations

The number of biologically relevant cooperator equilibria 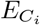 are independent of our bifurcation parameter *ρ* (because it does not appear in 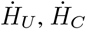, or 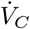) and depend instead on *r* and the burst size λ. The qualitative dynamics of Eq. (5) therefore vary with *r* and λ. Together with the insight that the number of biologically relevant infected equilibria of a single type depends on *r* relative to one, this suggests three parameter regimes:

1. host poor (*r* < 1): a single biologically relevant infected equilibrium for each viral type.
2. host rich & small burst size (*r* > 1, λ_*S*_ < λ < λ_*T*_): up to two biologically relevant infected equilibria for each viral type. Note that the two cooperator only equilibria emerge through a previously identified saddle node bifurcation at λ_*S*_ [37].
3. host rich & large burst size (*r* > 1, λ_*T*_ < λ): one biologically relevant infected equilibrium for cooperators, up to two for defectors.

In the host poor regime (*r* < 1), three transcritical bifurcations produce four dynamical regions, while in the host rich & large burst size regime (*r* > 1, λ_*T*_ < λ), seven bifurcations produce eight dynamical regions, and in the host rich & small burst size regime (*r* > 1, λ_*S*_ < λ < λ_*T*_), five bifurcations produce six dynamical regions, see Fig. 1. For full descriptions of each dynamical region, see appendix S2.

**Fig. 1.**
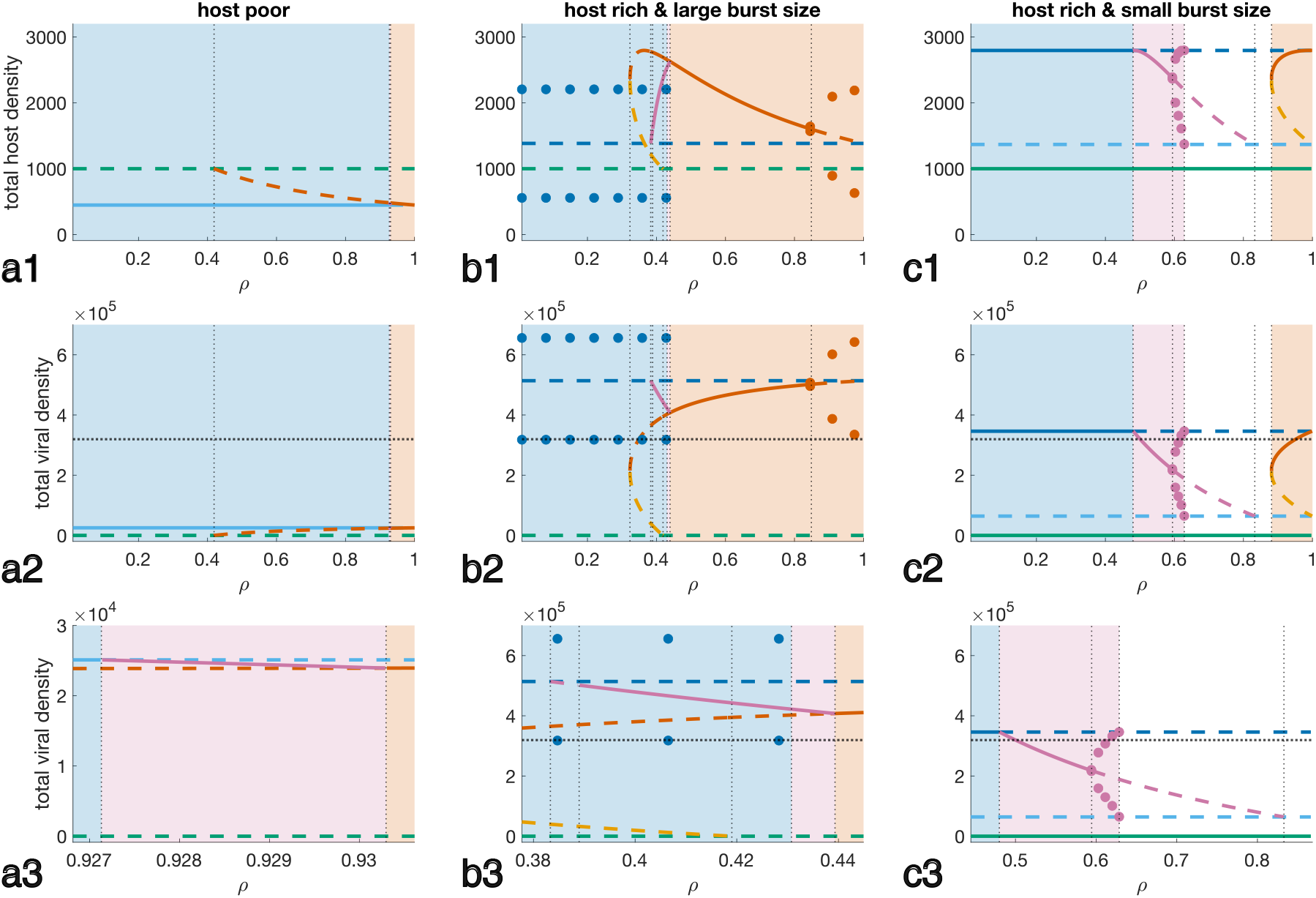
Bifurcation analysis reveals that environmental feedback maintains cooperation across three parameter regimes. Each column corresponds to a parameter regime; rows show (1) total host density (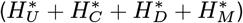), (2) total viral density (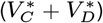), and (3) total viral density zoomed to the coexistence region. Line style indicates stability (solid, stable; dashed, unstable); color indicates equilibrium: green (*E*_*U*_), dark/light blue (*E*_*C*_), dark/light orange (*E*_*D*_), pink (*E*_*M*_). Circles indicate limit cycle amplitude. The dotted horizontal line in rows 2 and 3 marks *µ /µ*^2^, above which coinfection exceeds single infection. Background shading indicates long-term outcome after defectors are introduced to a cooperator population: blue, cooperators only; pink, coexistence; orange, defectors only; white, viral extinction. A) Host poor (*r* = 0.14, *λ* = 210): three bifurcations, four dynamical regions. B) Host rich, large burst size (*r* = 5, *λ* = 210): seven bifurcations, eight dynamical regions. C) Host rich, small burst size (*r* = 5, *λ* = 77): five bifurcations, six dynamical regions. **Parameters:** *r* = 0.14 [38]; *λ* = 210 [26]; *d* = 0.55 [26]; *µ* = 3.13 × 10^−6^, adapted from [26] (original value 9.939 × 10^−11^, increased to produce bifurcations at shorter timescales); *κ* = 1;*ξ* = *r/*1000.

Because lytic infection rapidly redirects host cellular machinery toward viral replication [35, 36], we assume that infected hosts do not compete with uninfected hosts for resources for replication (see Eq. (5a)). Consequently, in the two host rich regimes, uninfected hosts continue growing unimpeded as the infected class builds, driving total host density above the uninfected carrying capacity *r*/*ξ* (Fig. 1b1,c1). In the host poor regime (*r* < 1) this effect does not occur, and total host density at infected equilibria falls below *r*/*ξ* (Fig. 1a1).

### 3.4 Cooperation

The central question of our work is whether environmental feedback can provide an escape hatch for cooperation in viruses that face a prisoner’s dilemma in coinfection. In many regions, environmental feedback maintains cooperation: in blue shaded areas of Fig. 1 defectors go extinct but cooperators survive, while in pink shaded areas cooperators and defectors coexist either at stable equilibrium or through mixed limit cycles. However, environmental feedback is not always sufficient. In orange shaded areas of Fig. 1, cooperators go extinct while defectors survive. All three regimes follow the same general pattern: for small values of *ρ* cooperators can withstand the invasion of defectors and drive defectors extinct. Next follows a window in which cooperators and defectors co-exist. Finally, for sufficiently high *ρ*, defectors drive cooperators to extinction. The host rich & small burst size regime (*r* > 1, λ_*S*_ < λ < λ_*T*_) allows an additional outcome: in the white shaded areas of Fig. 1, the introduction of defectors drives both types extinct; the implications of defector driven viral extinction are explored further below.

Environmental feedback maintains cooperation because it allows each type to benefit (cooperators) or suffer (defectors) from growth rate differences during single infection, in the absence of any interaction. In Eq. (5), the rates of single infection versus coinfection vary with viral and host density: single infections occur at rate *µ H*_*U*_ (*V*_*C*_ + *V*_*D*_) while double infections occur at rate *µ*^2^*H*_*U*_ (*V*_*C*_ + *V*_*D*_)^2^. It follows that coinfection is more common than single infection when *V*_*C*_ + *V*_*D*_ > 1/*µ*, and vice versa. The horizontal dotted line at 1/*µ* in Fig. 1 differentiates the two cases. In the host poor regime (*r* < 1), coinfection accounts for around 7% of infection events at the cooperator equilibrium. In this case a type’s fitness is determined primarily by its single-infection replication rate, and defectors cannot invade until their replication rate *ρ* approaches that of cooperators (*ρ* > *ρT*_*MC*_ *≈* 0.9271). In the two host rich regimes, viral densities are higher and coinfection has more influence on each type’s fitness. In both regimes, coinfection is more common than single infection in cooperator populations (large burst size: coinfections fluctuate between 50-67% of infections; small burst size: coinfection accounts for 52% of infections) . However, this does not guarantee the spread of defectors, which still cannot invade until defector replication in single infection is sufficiently high (host rich & large burst size *ρ* > *ρT*_*MD*_ *≈* 0.4393; host rich & small burst size 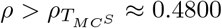). In summary, even when coinfection is common, single infections that penalize defectors and advantage cooperators still occur and influence dynamics.

The balance between single and coinfection and feedback through viral densities is well illustrated through mixed limit cycles of cooperators and defectors in the host rich & small burst size regime (pink circles, Fig. 1**c1-3**; *r* > 1, λ_*S*_ < λ < λ_*T*_). In Fig. 2, cooperator-rich populations (blue) expand in density, increasing the relative rate of coinfection as each trajectory approaches the dashed line. Increased coinfection favors defectors (transition to yellow), which increase in density at the expense of cooperators. High defector frequencies make coinfecting with a cooperator less likely, decreasing defector replication until defector densities drop. Coinfection is rare far from the dashed line, allowing cooperators to benefit from their fitness advantage in single infection and rebound (transition back to blue). These mixed limit cycles recapitulate previously observed oscillations in the defector-to-cooperator ratio, termed the von Magnus effect [8, 9, 30, 39], and support prior suggestions that environmental feedback drives the effect [40].

**Fig. 2.**
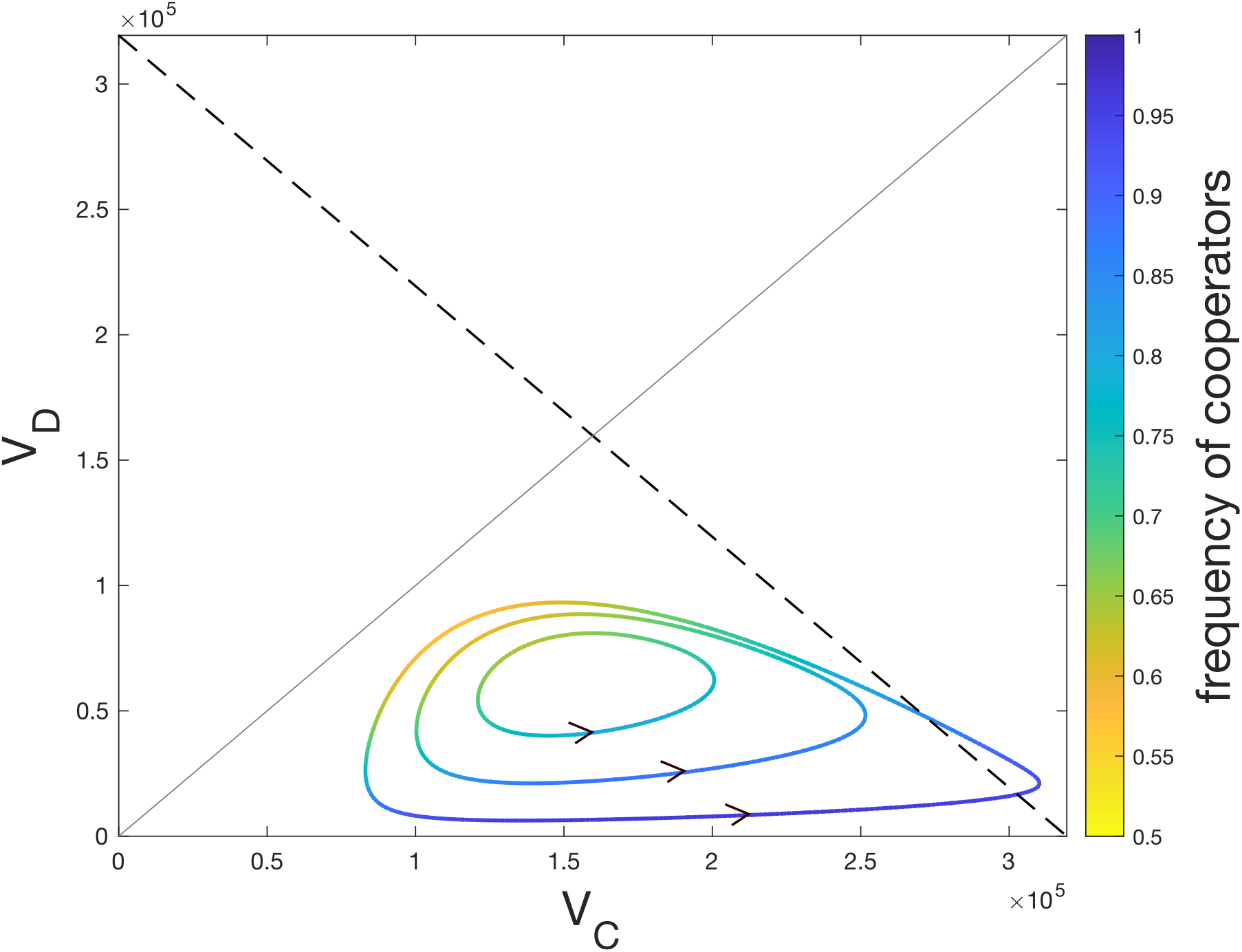
Stable mixed limit cycles in the host rich & small burst size regime (*r >* 1, *λ*_*S*_ *< λ < λ*_*T*_). For *ρ > ρH*_*M*_ a limit cycle appears and grows with *ρ* until it is lost at *ρ*_*HC*_ . On each trajectory, color indicates the frequency of cooperation at that time point, *V*_*C*_ */*(*V*_*C*_ + *V*_*D*_); this can also be inferred by comparing to the solid line *V*_*D*_ = *V*_*C*_ . Above dotted line *V*_*C*_ + *V*_*D*_ = 1*/µ* coinfection is more common, while below the line single infection is more common. **Parameters:** as in Fig. 1; *r* = 5, *λ* = 77, *ρ* ∈ {0.6, 0.61, 0.62}.

### 3.5 Implications for viral applications

#### 3.5.1 Defectors hinder the use of phages as an alternative antibiotic

In the host rich & large burst size regime (*r* > 1, λ_*T*_ < λ), cooperator populations exhibit limit cycle behavior, see Fig. 1b1-3. The *H*_*U*_ component cycles close to zero, see Fig. 3a, meaning that the uninfected hosts are periodically close to extinction. Stochastic effects are important in populations with small numbers [41, 42] so small perturbations, either through natural stochastic variation in host densities, or through a well timed intervention, could easily result in extinction of the host population [37]. These dynamics support the proposed application of phages as an alternate antibiotic to eliminate host bacterial populations.

**Fig. 3.**
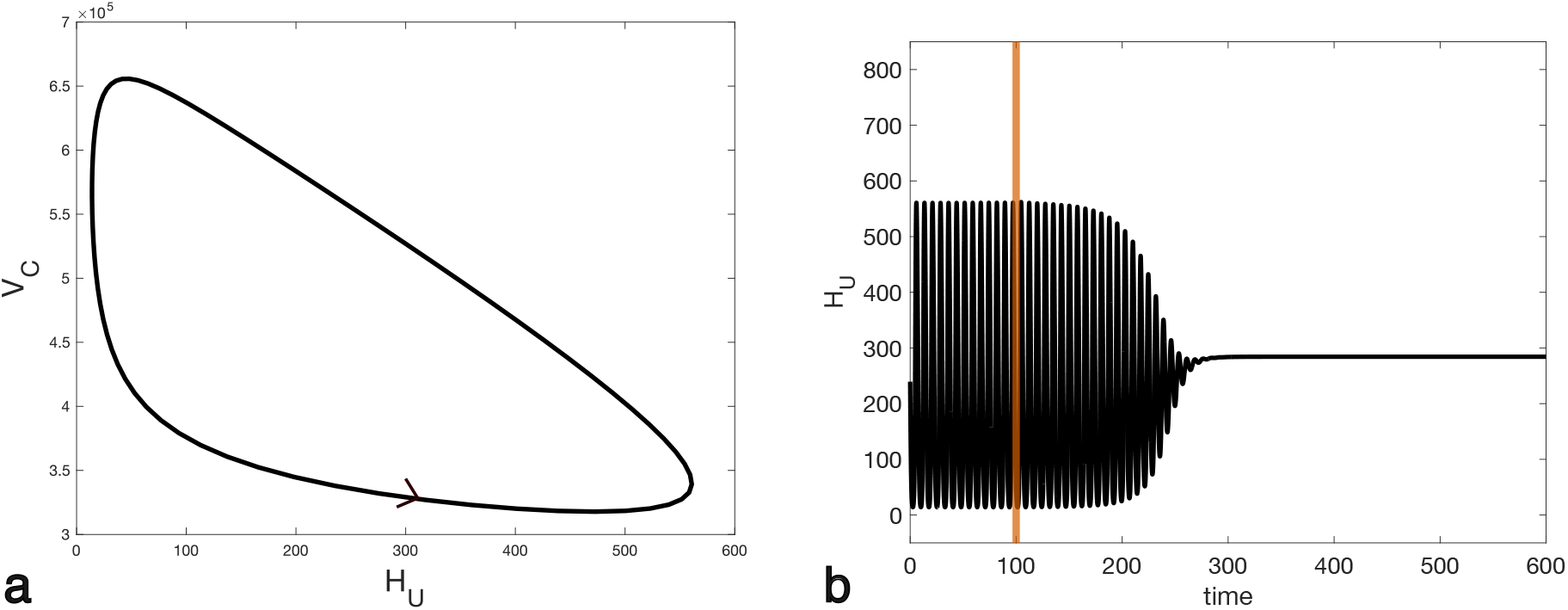
Defectors eliminate the risk of host extinction in the host rich & large burst size regime (*r >* 1, *λ*_*T*_ *< λ*). Panel **a** shows a projection of the cooperator limit cycle in *H*_*U*_, *V*_*C*_ phase space. Uninfected hosts periodically have densities close to zero. Panel **b** shows a time series: at the vertical orange line, defectors are introduced into a cycling cooperator only population. The system moves to the defector equilibrium, where host densities are high and stable. **Parameters:** as in Fig. 1; *r* = 5, *λ* = 210, *ρ* = 0.6.

However, for *ρT*_*MD*_ < *ρ* < *ρH*_*D*_ (orange region of Fig. 1b1-3) defectors can invade cooperators and drive them extinct, resulting in a population of defectors at stable equilibrium, where the risk of stochastic extinction of the host is minimal, see Fig. 3b. As *ρ* increases, defectors eventually exhibit limit cycle behavior, but the host is still less prone to stochastic extinction until *ρ* is close to one. In interpretation, the introduction of defectors compromises the use of phages to eliminate bacteria.

#### 3.5.2 Introduction of defectors lowers viral densities and drives viral extinction

Across all three regimes, successful invasion of defectors into a population of cooperators results in lower total viral densities, though this reduction wanes as *ρ* increases towards one. Further, the introduction of defectors guarantees viral extinction for 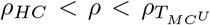 (white region of Fig. 1c1-3). In this regime defectors cannot establish a successful infection in the absence of cooperators, and further cooperator only populations are bi-stable between the stable cooperator equilibrium *E*_*C*_ and *E*_*U*_ . In Fig. 4a, coinfection is common at the cooperator only equilibrium (trajectory begins above the dashed threshold line). After introduction, rare defectors coinfect with cooperators and rapidly increase in density at the expense of cooperators until defectors are common (yellow portion of trajectory, defector frequency *≈* 0.5). Now coinfections are increasingly between two defectors rather than a defector and cooperator. Defectors lose their advantage and defector densities drop (transition from yellow to blue). Coinfection is rare, accounting for 10% or less of total infection events once defectors drop below frequencies of 0.25, and we expect defector and cooperator dynamics to be largely decoupled. Defector replication is too low for defectors to sustain themselves (*R*_0_ < 1 for defectors and no defector equilibrium), so the defector type is lost, and at this point cooperators are in the basin of attraction for *E*_*U*_, see Fig. 4b, so cooperators are lost as well.

**Fig. 4.**
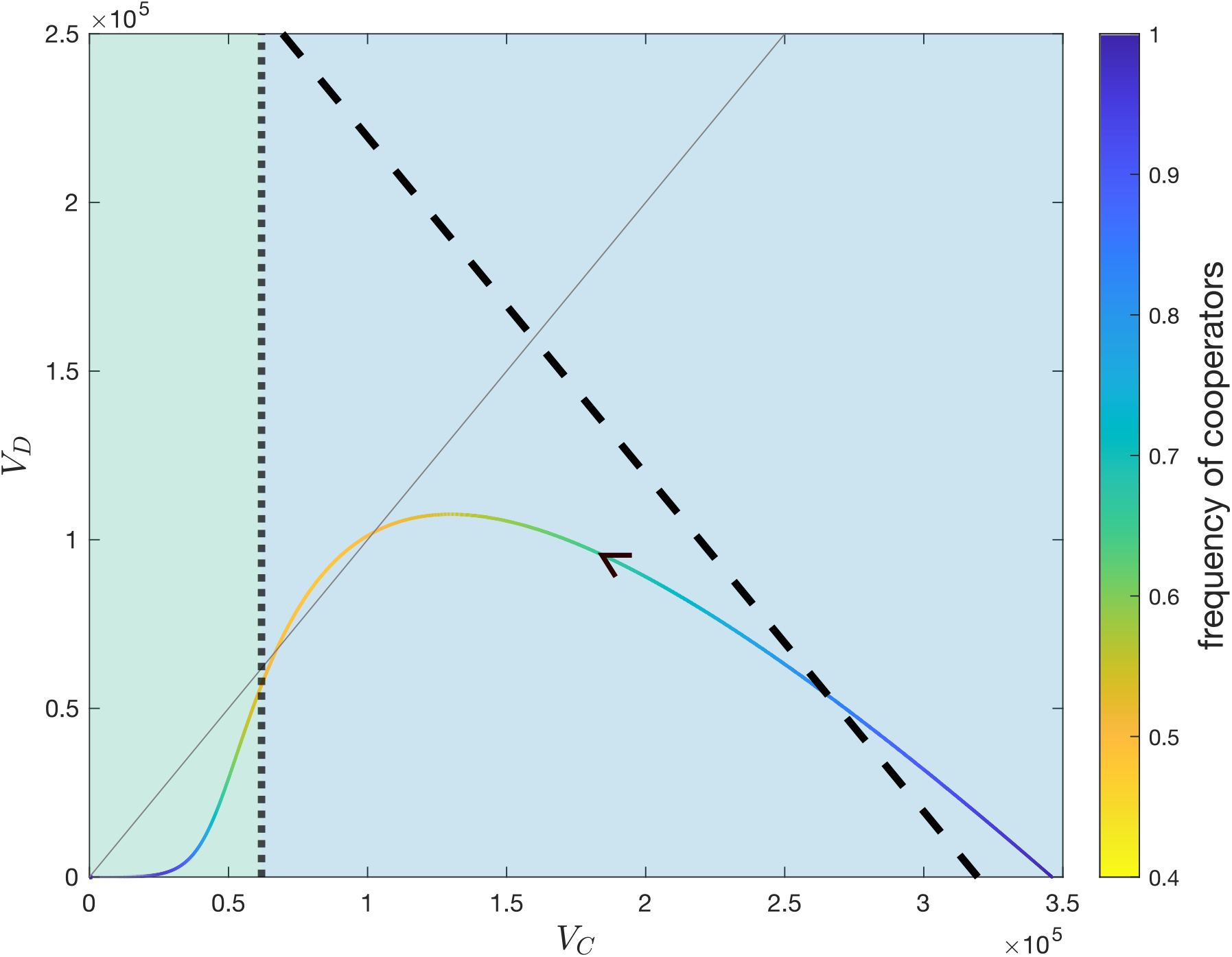
Introduction of defectors drives viral extinction in the host rich & small burst size regime (*r >* 1, *λ*_*S*_ *< λ < λ*_*T*_). The solid, colored line shows a population trajectory after a defector is introduced into a cooperator population, projected into *V*_*C*_, *V*_*D*_ phase space. Color corresponds to the frequency of cooperators at a given point in time, *V*_*C*_ */*(*V*_*C*_ + *V*_*D*_); this can also be inferred by referencing the solid line showing *V*_*D*_ = *V*_*C*_ . Coinfection is more common than single infection above the dashed line *V*_*C*_ + *V*_*D*_ = 1*/µ*, and vice versa below the line. Background shading indicates the basin of attraction for a given point on the trajectory with no defectors. Points in the blue region (right of dashed line) are drawn to 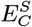, while points in the green region (left of dashed line) are drawn towards *E*_*U*_ . **Parameters:** as in Fig. 1; *r* = 5, *λ* = 77, *ρ* = 0.65.

## 4 Discussion

We show that environmental feedback can maintain cooperation in viruses. Here, environmental feedback is defined as changes in population composition that alter the environment (viral and host densities) and feed back to affect the fitness of each type (by altering the frequency of single infection versus coinfections). Environmental feedback is a robust mechanism that maintains cooperation across all ecological scenarios considered. In some regions, cooperators drive defectors to extinction (blue backgrounds in Fig. 1), while in others cooperators and defectors coexist (pink backgrounds in Fig. 1), either at a stable equilibrium or in a mixed limit cycle analogous to previously observed oscillations of cooperator and defector density [8, 30, 39]. However, environmental feedback is not always sufficient to overcome the prisoner’s dilemma viruses face in coinfection; if defectors gain a sufficient fitness advantage over cooperators, defectors can still drive cooperators to extinction (orange backgrounds in Fig. 1).

Our model was inspired by the dynamics of bacteriophages and their bacterial hosts. Because lytic infection rapidly redirects host cellular machinery toward viral replication [35, 36], in our model infected hosts do not compete with uninfected hosts for resources. As a result, uninfected hosts continue to grow even as infected hosts accumulate, raising the total host density above the uninfected host carrying capacity when host growth rates are high (*r* > 1; Fig. 1b1,c1). This increases viral densities and the frequency of coinfection, conditions that favor defectors. Yet even here environmental feedback maintains cooperation. We anticipate that our results should hold generally when the environment enables changes in population composition to feed back and affect the relative fitness of cooperators and defectors.

Environmental feedback shares structural features with two previously identified mechanisms that maintain cooperation. In parallel to group selection, here within-cell dynamics create a between-cell advantage for cooperators because cooperator-dominated cells are more productive than defector-dominated cells [20]. At the population level, the spread of defectors reduces overall viral density and increases the frequency of single infections that advantage cooperators, similar to work incorporating ecological dynamics where the spread of defectors leads to smaller groups that favor cooperators [21]. Both previous mechanisms operate within the framework of the replicator equation, which assumes that all fitness differences between types arise from their interactions [43], and cooperation is maintained by ensuring that cooperators interact preferentially [44].

However, environmental feedback is fundamentally distinct because it incorporates fitness effects that are independent of interactions entirely. Though viruses experience the prisoner’s dilemma during coinfection, cooperators have higher rates of replication than defectors when each type infects a cell alone, a dynamic not captured within the replicator framework. Environmental feedback maintains cooperation by balancing coinfections, where defectors are advantaged, and single infections, where cooperators are advantaged. Even when coinfection is more common, the cooperator advantage in single infections maintains cooperation as long as defectors’ replication advantage is not too high.

Environmental feedback offers viruses an ecological escape from the mutual defection predicted by the prisoner’s dilemma. Previous work has identified evolutionary escapes from the prisoner’s dilemma. When infecting cells alone, phage *ϕ*_6_ evolves an efficient cooperator phenotype that plays a snowdrift game (*γ* > *α* > *β* > *δ* in Eq. (1)), rather than a prisoner’s dilemma, with defectors [45]. In addition, nonlinear trade-offs in public goods production can shift the game from prisoner’s dilemma to snowdrift even under coinfection [46]. In either case, mutual defection is no longer dominant and long-term coexistence is predicted. The evolutionary escape requires payoffs to change, altering the interaction between cooperators and defectors, while the ecological escape does not. Cooperators always play a prisoner’s dilemma in coinfection, but environmental feedback determines how often coinfection occurs. Our results highlight the need to consider the ecology of a system, beyond interactions between types, when predicting the fate of cooperation [22].

Phage therapy proposes the use of bacteriophages as an alternative to antibiotics for eliminating pathogenic bacterial populations [47]. We show that defectors can undermine this application by disrupting the viral dynamics that drive host extinction. In cooperator populations, limit cycle oscillations drive uninfected host densities close to zero, where stochastic effects [41, 42] or well-timed interventions can cause bacterial extinction [37]. The introduction of defectors eliminates this risk: the system moves instead to a stable equilibrium at which uninfected bacterial densities remain high. Viral defectors are known to arise spontaneously, especially when the density of phages is high relative to bacteria and many phages coinfect [10, 27]. Screening for defector evolution during treatment development and administration is critical for preserving therapeutic efficacy. However, some viral defectors, such as those generated by point mutations, are difficult to detect [25]. In light of this, a good precaution is to ensure that phage are cultured in conditions where coinfection is rare. In addition, administering low doses of phage could prevent coinfection and delay the evolution of defectors that hamper the treatment, though this is at odds with current dosing recommendations which suggest administering high density of phage relative to bacteria [48] so that each bacterial cell is swiftly infected.

Viral defectors have been harnessed as antiviral therapies across multiple viral species [49–57]. Most current therapeutic defectors are incapable of replicating without a cooperator [58]. Our results suggest a wider range of defectors should be considered as therapeutic candidates. In our model, partial defectors capable of replicating independently can drive total extinction (white region, Fig. 1, Fig. 4). Beyond this, if a reduction in viral load, rather than total viral extinction, is a reasonable therapeutic goal [49], an even wider range of partial defectors might be considered, as in our results the introduction of defectors reduces viral densities as long as defector replication in single infection is lower than that of cooperators.

## 5 Acknowledgments

We thank Daniel Coombs, Kayla King, Asher Leeks, and Sally Otto, as well as Luis Izquierdo, Segismundo Izquierdo, and George Berry for helpful discussion. The authors acknowledge funding by the Natural Sciences and Engineering Research Council of Canada, Discovery Grant RGPIN-2021-02608 to C.H.

