## Supplementary material for "Environmental feedback maintains cooperation in viruses"

### S1 Stability

The Jacobian of Eq. (5) has the form below, where $V = V_C + V_D$.

$$\begin{pmatrix} r - 2H_U\xi - \mu V - \mu^2V^2 & 0 & 0 & 0 & -H_U(\mu + 2\mu^2 V) & -H_U(\mu + 2\mu^2 V) \\ V_C(\mu + \mu^2 V_C) & -d & 0 & 0 & H_U(\mu + 2\mu^2 V_C) & 0 \\ V_D(\mu + \mu^2 V_D) & 0 & -d & 0 & 0 & H_U(\mu + 2\mu^2 V_D) \\ 2\mu^2 V_C V_D & 0 & 0 & -d & 2\mu^2 H_U V_D & 2\mu^2 H_U V_C \\ 0 & 2d\lambda & 0 & 0 & -d\kappa & 0 \\ 0 & 0 & 2d\lambda\rho & d\lambda(1+\rho) & 0 & -d\kappa \end{pmatrix}.$$

The Jacobian evaluated at $E_0$ has eigenvalues $[0, r, -d, -d\kappa, -d, -d\kappa]$, and thus $E_0$ is unstable for $r > 0$. The Jacobian evaluated at $E_U$ has eigenvalues

$$\lambda_1 = -d$$
$$\lambda_2 = -r$$
$$\lambda_{3,4} = \frac{-d(1+\kappa) \pm \xi\sqrt{\frac{d^2\xi(\kappa-1)^2 + 8d\lambda\mu\, r}{\xi^3}}}{2}$$
$$\lambda_{5,6} = \frac{-d(1+\kappa) \pm \xi\sqrt{\frac{d^2\xi(\kappa-1)^2 + 8d\lambda\mu\, r\rho}{\xi^3}}}{2}$$

Eigenvalues $\lambda_3$ and $\lambda_5$ can be positive. If either is positive, $E_U$ is unstable. The conditions for which $\lambda_3$ and $\lambda_5$ are positive correspond to $R_0 > 1$ for cooperators and defectors, respectively. Eigenvalue $\lambda_3$ is positive for $\lambda > (d\xi\kappa)/(2\mu\, r)$. To see this, consider

$$\xi\sqrt{\frac{d^2\xi(\kappa-1)^2 + 8d\lambda\mu\, r}{\xi^3}} > d(1+\kappa).$$

Note that both sides are positive. After squaring and rearranging, we obtain

$$8\lambda\mu\, r\, d > 4d^2\xi\kappa$$

which yields the above threshold for $\lambda$. Following similar algebraic steps, we show that eigenvalue $\lambda_5$ is positive for $\rho > (d\xi\kappa)/(2\lambda\mu\, r)$.

### S2 Model dynamics

#### S2.1 Host poor regime ($r < 1$)

There are three transcritical bifurcations: first between $E_U$ and $E_D$ at $\rho_{T_{DU}} = (d\kappa\xi) / (2\lambda\mu r) \approx 0.4190$, then between $E_M$ and $E_C$, which exchange stability, at $\rho_{T_{MC}} \approx 0.9271$, and finally between $E_M$ and $E_D$, which again exchange stability, at $\rho_{T_{MD}} \approx 0.9302$. For the exact form of transcritical bifurcations involving $E_M$, see appendix S3. These bifurcations lead to four dynamical regions, described below and represented graphically in Fig. S1.

a **Cooperators only** ($\rho < \rho_{T_{DU}}$): The cooperator equilibrium $E_C$ and uninfected equilibrium $E_U$ are biologically relevant, but only $E_C$ is stable. Specifically, $R_0 > 1$ for cooperators, which can establish an infection from arbitrarily small densities in the absence of defectors. In contrast, the defector and mixed equilibria are not biologically relevant, so defectors invariably go extinct. See Fig. S1A.

b **Cooperators exclude defectors** ($\rho_{T_{DU}} < \rho < \rho_{T_{MC}}$): The defector equilibrium $E_D$ becomes biologically relevant in a transcritical bifurcation with the uninfected equilibrium $E_U$ at $\rho_{T_{DU}}$. Now $R_0 > 1$ for defectors and the type can establish an infection from arbitrarily small densities in the absence of cooperators. However, $E_C$ remains the only attractor: cooperators can invade defectors, driving them extinct, while defectors cannot invade cooperators. See Fig. S1B.

c **Coexistence** ($\rho_{T_{MC}} < \rho < \rho_{T_{MD}}$): At $\rho_{T_{MC}}$ the cooperator equilibrium $E_C$ loses stability through a transcritical bifurcation with the mixed equilibrium $E_M$, which gains stability. Now the mixed equilibrium $E_M$ is the sole stable equilibrium: cooperators and defectors can each invade the other type, leading to coexistence. See Fig. S1C.

d **Defectors exclude cooperators** ($\rho_{T_{MD}} < \rho < 1$): At $\rho_{T_{MD}}$ the defector only equilibrium $E_D$ gains stability and is the only stable attractor. Defectors can invade cooperators and drive them to extinction, but cooperators cannot invade defectors. See Fig. S1D.

#### S2.2 Host rich & large burst size regime ($r > 1$, $\lambda > \lambda_T$)

There are seven bifurcations: first a stable/unstable pair of defector equilibria appear in a saddle node bifurcation at $\rho_{S_D} = d\kappa\xi \left(-2 - 9r + 2\sqrt{(1+3r)^3}\right) / \left(2\mu r^2(1+4r)\lambda\right) \approx 0.3232$. Referencing Fig. 1b, we denote the top and bottom branches $E_{D^S}$ and $E_{D^U}$, respectively. This is followed by a transcritical bifurcation between the mixed and cooperator only equilibria at $\rho_{T_{MC}} \approx 0.3833$. Shortly after the mixed equilibrium gains stability at $\rho_M = 0.3889$. Next is a transcritical bifurcation between the uninfected and a defector only equilibria at $\rho_{T_{DU}} = (d\kappa\xi) / (2\lambda\mu r) \approx 0.4190$, after which the cooperator only limit cycle loses stability at $\rho_{LC} = 0.4307$

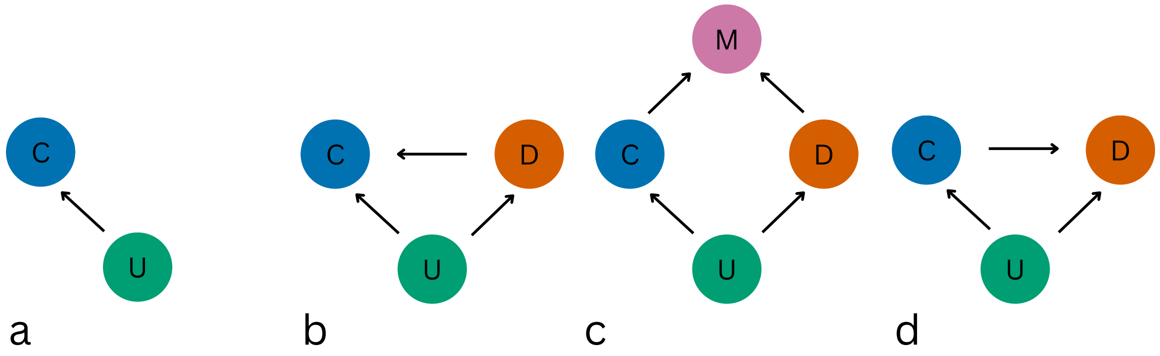


**Fig. S1.** Long term population composition for the host poor regime ($r < 1$). Circles correspond to equilibria: U, uninfected; C, cooperator only; D, defector only; M, mixed. Arrows indicate that viral invasion can lead from one equilibria to the other. In **A**, only cooperators can invade and establish an infection ($R_0 > 1$ for cooperators). In **B**, both cooperators and defectors can invade ($R_0 > 1$ for both types). In addition, cooperators can invade defectors and drive them to extinction. In **C** both cooperators and defectors can invade the other type, resulting in a mixed state. In **D**, defectors can invade cooperators, but cooperators cannot invade defectors.

as its basin of attraction shrinks to cooperator-only trajectories. Next is another transcritical bifurcation, this time between the mixed and a defector only equilibria at $\rho_{T_{MD}} \approx 0.4393$. Finally, there is a Hopf-bifurcation as the defector only equilibrium loses stability at $\rho_{H_D} \approx 0.8490$, resulting in defector only limit cycles that have been confirmed numerically. These bifurcations lead to eight dynamical regimes, described in greater detail below and represented graphically in Fig. S2.

a **Cooperator limit cycles** ($\rho < \rho_{S_D}$): The uninfected and cooperator equilibria, $E_C$ and $E_U$, are biologically relevant but both are unstable; cooperator limit cycles are anticipated and have been confirmed numerically. No defector or mixed equilibria are biologically relevant, so defectors invariably go extinct. Reference Fig. S2A.

b **Cooperators exclude defectors** ($\rho_{S_D} < \rho < \rho_{T_{MC}}$): At $\rho_{S_D}$ a pair of defector equilibria appear through a saddle node bifurcation; referencing Fig. 1b, we denote the top and bottom branch $E_D^S$ and $E_D^U$, respectively, as in the absence of cooperators the branches are respectively stable and unstable. In the absence of cooperators, defector dynamics are bi-stable between $E_U$ and $E_D^S$; trajectories in the basin of attraction for $E_D^S$ will establish a successful infection, while trajectories in the basin of attraction for $E_U$ lose all virus. Defectors cannot invade cooperators, though cooperators can invade defectors and drive them extinct. Reference Fig. S2B.

c **Cooperators exclude defectors II** ($\rho_{T_{MC}} < \rho < \rho_M$): At $\rho_{T_{MC}}$ the mixed equilibrium $E_M$ becomes biologically relevant but is unstable. Stable cooperator limit cycles remain the only attractor, and the dynamics are unchanged from the previous region. Reference Fig. S2C.

d **Bistability between cooperator limit cycles and coexistence** $(\rho_M < \rho < \rho_{T_{DU}})$ : At $\rho_M$ the mixed $E_M$ gains stability. Dynamics are bi-stable between the mixed equilibrium $E_M$ and cooperator limit cycles. More specifically, cooperators can invade defectors, leading to coexistence, but defectors cannot invade cooperators. Reference Fig. S2D.

e **Bistability between cooperator limit cycles and coexistence II** $(\rho_{T_{DU}} < \rho < \rho_{T_{LC}})$: At $\rho_{T_{DU}}$ the defector equilibrium $E_D^U$ becomes biologically irrelevant through a transcritical bifurcation with $E_U$. Now $R_0 > 1$ for a defector-only population, and the type can establish an infection from arbitrarily small densities in the absence of cooperators. All other dynamics are identical to the previous region. Reference Fig. S2E.

f **Coexistence** $(\rho_{LC} < \rho < \rho_{T_{MD}})$: The cooperator limit cycle loses stability at $\rho_{LC}$. Both cooperators and defectors can invade the other type, leading to coexistence. Reference Fig. S2F.

g **Defectors exclude cooperators** $(\rho_{T_{MD}} < \rho < \rho_{H_D})$: At $\rho_{T_{MD}}$ the mixed equilibrium $E_M$ transfers stability to $E_D^S$ through a transcritical bifurcation. The defector equilibrium, $E_D^S$ is now the sole attractor. Cooperators cannot invade defectors, but defectors can invade cooperators and drive them extinct. Reference Fig. S2G.

h **Defector limit cycles** $(\rho_{H_D} < \rho < 1)$: At $\rho_{H_D}$ the defector equilibrium $E_{D^S}$ loses stability in a Hopf bifurcation; no stable equilibria remain, and defector limit cycles are anticipated and have been confirmed numerically. Cooperators cannot invade defectors, but defectors can invade cooperators and drive them extinct. Reference Fig. S2H.

#### S2.3 Host rich & small burst size regime ($r > 1$, $\lambda_S < \lambda < \lambda_T$)

There are five bifurcations: first a transcritical bifurcation between $E_M$ and $E_C^S$ at $\rho_{T_{MC^S}} \approx 0.4800$. Next, $E_M$ loses stability through a Hopf-bifurcation at $\rho_{H_M} \approx 0.5944$; mixed limit cycles have been confirmed numerically. Next the mixed limit cycle loses stability through an apparent homoclinic bifurcation [1] at $\rho_{HC} \approx 0.6286$ as the limit cycle intersects $E_C^U$, see Fig. S3. Next, there is a transcritical bifurcation between $E_M$ and $E_C^U$ at $\rho_{T_{MC^U}} \approx 0.8330$, and finally a stable/unstable pair of defector equilibria appear in a saddle node bifurcation at $\rho_{S_D} = d\kappa\xi\left(-2-9r+2\sqrt{(1+3r)^3}\right) / \left(2\mu r^2(1+4r)\lambda\right) \approx 0.8816$. These bifurcations lead to six dynamical regimes, described in greater detail below and illustrated in Fig. S4.

a **Bistability between $E_C^S$ and $E_U$** $(\rho < \rho_{T_{MC^S}})$: Two cooperator only equilibria are biologically relevant; referencing Fig. 1c we denote the top and bottom equilibria $E_C^U$ and $E_C^S$, respectively. Both $E_C^S$ and $E_U$ are stable, so dynamics are bistable. Trajectories within the basin of attraction for $E_C^U$ will establish a

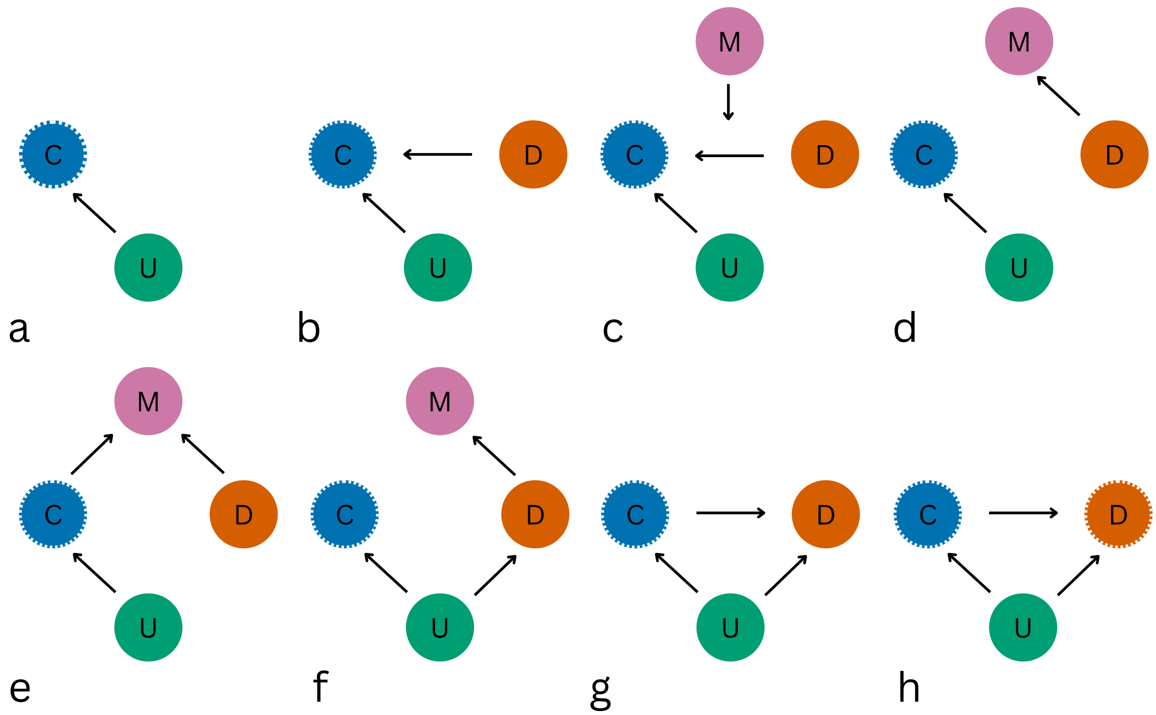


**Fig. S2.** Long term population composition for the host rich & large burst size regime ($r > 1$, $\lambda > \lambda_T$). Circles with no border correspond to equilibria and circles with dashed borders correspond to stable limit cycles: U, uninfected; C, cooperator only; D, defector only; M, mixed. Arrows indicate that viral invasion can lead from one composition to the other. In **A**, cooperators can invade ($R_0 > 1$ for cooperators) leading to cooperator limit cycles. In **B**, defectors can establish from a sufficiently high initial density ($R_0 < 1$ for defectors), leading to a defector only state. Cooperators can invade defectors and drive them to extinction, leading to cooperator limit cycles. In **C**, a population at precisely the mixed equilibrium would remain in a mixed state, but overall the long term population dynamics are equivalent to B. In **D** cooperators can invade defectors, leading to a mixed state. In **E** defectors can invade cooperators, leading to a mixed state. The dynamics of **F**, are identical to E but now defectors can invade an uninfected host population ($R_0 > 1$ for defectors), leading to a defector only state. In **G** Defectors can invade cooperators and drive them to extinction. In **H** defectors experience limit cycle dynamics.

successful infection, while trajectories within the basin of attraction for $E_U$ lose all virus. Neither the mixed defector only or mixed equilibrium are stable, so defectors invariably go extinct. See Fig. S4A.

b **Coexistence** ($\rho_{T_{MCS}} < \rho < \rho_{H_M}$): The mixed equilibrium $E_M$ appears in a transcritical bifurcation with $E_{C^S}$, which loses stability, at $\rho_{T_{MCS}}$. Dynamics are bi-stable between the uninfected equilibrium, $E_U$, and the mixed equilibrium, $E_M$. Specifically, defectors can invade cooperators, leading to stable co-existence of both types. Reference Fig. S4B.

c **Coexistence through limit cycles** ($\rho_{H_M} < \rho < \rho_{HC}$): At $\rho_{H_M}$ the mixed equilibrium $E_M$ loses stability in a Hopf bifurcation; mixed limit cycles have been confirmed numerically. Dynamics are bi-stable between the $E_U$ and mixed limit cycles: defectors can invade cooperators, though now the two types coexist through the turn of a limit cycle. Reference Fig. S4C.

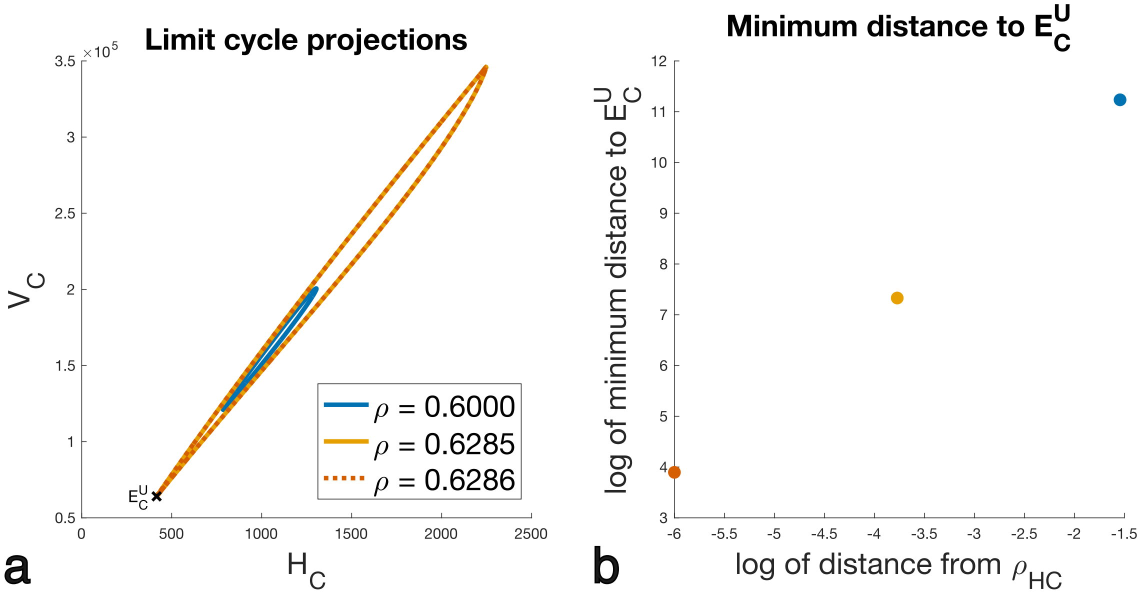


**Fig. S3.** Evidence for a homoclinic bifurcation at $\rho_{HC} \approx 0.62864$. Colors are shared between panels. **a** Projections of the mixed limit cycle onto the $(H_C, V_C)$ plane; As $\rho \to \rho_{HC}$, the limit cycle grows and approaches $E_C^U$. Recall that all cooperator equilibria are independent of $\rho$, so $E_C^U$ does not move as $\rho$ varies. **b** Logarithm of minimum Euclidean distance from each limit cycle to $E_C^U$, plotted against log-distance from $\rho_{HC}$ (right = far from $\rho_{HC}$, left = close). The monotonic decrease toward zero is consistent with the limit cycle colliding with $E_C^U$ at $\rho_{HC}$. Parameters as in Fig. 1; $r = 5$, $\lambda = 77$.

d **Extinction** ($\rho_{HC} < \rho < \rho_{T_{MC^U}}$): At $\rho_{HC}$ the mixed limit cycles are lost in a likely homoclinic bifurcation as the mixed limit cycle collides with the cooperator equilibrium $E_C^U$, see Fig. S3. Now only the uninfected equilibrium $E_U$ is stable: the introduction of defectors into a cooperator population drives the entire viral population to extinction. Reference Fig. S4D.

e **Extinction II** ($\rho_{T_{MC^U}} < \rho < \rho_{S_D}$): At $\rho_{T_{MC^U}}$ the mixed equilibrium $E_M$ disappears in a transcritical bifurcation with $E_{C^U}$ . Dynamics are qualitatively identical to region D. Reference Fig. S4E.

f **Defectors exclude cooperators** ($\rho_{S_D} < \rho < 1$): At $\rho_{S_D}$ a stable/unstable pair of defector equilibria appear in a saddle node bifurcation. Referencing Fig. 1c, we denote the top and bottom branches $E_{D^S}$ and $E_{D^U}$, respectively. Dynamics are bi-stable between the $E_U$ and $E_D^S$. Cooperators cannot invade a population of defectors, while defectors can invade cooperators and drive them to extinction. Reference Fig. S4F.

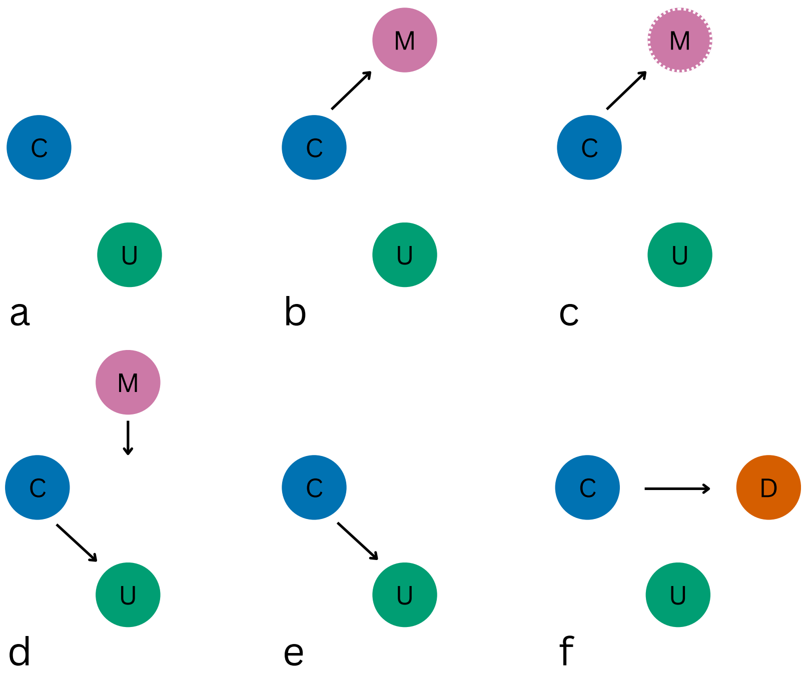


**Fig. S4.** Long term population composition for the host rich & small burst size regime ($r > 1$, $\lambda_S < \lambda < \lambda_T$). Circles with no border correspond to equilibria and circles with dashed borders correspond to stable limit cycles: U, uninfected; C, cooperator only; D, defector only; M, mixed. Arrows indicate that viral invasion can lead from one composition to the other. In **A**, cooperators can establish from sufficiently high initial densities ($R_0 < 1$)), leading to a cooperator only state. In the next panels, defectors can invade cooperators, leading to a mixed state, either at stable equilibrium **B** or in a mixed limit cycle **C**. In **D**/**E**, defector invasion of the cooperator only state drives the entire viral population to extinction. In **F**, defectors can establish from sufficiently high initial densities ($R_0 < 1$ for defectors), leading to a defector only state. In addition, defectors can invade cooperators and drive them to extinction.

### S3 Transcritical bifurcations of the mixed equilibrium

To find the exact form of transcritical bifurcations involving the mixed equilibrium, $E_M$, we are interested in collisions between the mixed equilibrium and either cooperator or defector equilibria, $E_{C_i}$ or $E_{D_i}$. Conveniently, $E_M$ has a form that is analytically tractable, though it is not readable.

To find intersections between $E_M$ and $E_{C_i}$, we seek $\rho^*_{C_i}$ such that the mixed equilibrium evaluated at $\rho^*_{C_i}$ has the form

$$E_M|_{\rho=\rho^*_{C_i}} = [H^*_U > 0, H^*_C > 0, H^*_D = 0, H^*_M = 0, P^*_C > 0, P^*_D = 0].$$

Setting the $H^*_D$ component of $E_M$ equal to 0, we find candidate $\rho^*_{C_i}$ which are the roots of the cubic polynomial

$$f(\rho) = d\kappa\xi\rho^3 - 2\lambda\mu\, r\rho^2 - 2\lambda\mu\rho + 2\lambda\mu. \quad \text{[S1]}$$

We confirm that $E_M$ evaluated at the roots of this cubic polynomial has the desired form, and further that in each parameter regime, transcritical bifurcations between $E_M$ and $E_{C_i}$ correspond to one of the roots.

Similarly, to find intersections between $E_M$ and $E_{D_i}$, we seek $\rho^*_{D_i}$ such that the mixed equilbrium evaluated at $\rho^*_{D_i}$ has the form

$$E_M|_{\rho=\rho^*_{D_i}} = [H^*_U > 0, H^*_C = 0, H^*_D > 0, H^*_M = 0, P^*_C = 0, P^*_D > 0].$$

Setting the $H^*_C$ component of $E_M$ equal to zero we find two candidate $\rho^*_{D_i}$, $\frac{\lambda\,\mu \pm \sqrt{\lambda\,\mu\,(\lambda\,\mu - 2\,d\,\kappa\,\xi + 4\,\lambda\,\mu\,r)}}{d\,\kappa\,\xi - 2\,\lambda\,\mu\,r}$, and confirm that $E_M$ evaluated at these $\rho^*_{D_i}$ has the correct form, and that in each regime transcritical bifurcations between $E_M$ and $E_{D_i}$ correspond to one of these roots.
